# Distributed neurophysiological dynamics link perception, action, and language in schizophrenia

**DOI:** 10.64898/2026.02.17.706352

**Authors:** Hsi T. Wei, Dominic Boutet, Rukun Dou, Jessica Ahrens, Nadia Zeramdini, Alban Voppel, Fernando Gonzales-Aste, Fatme Abboud, Sylvain Baillet, Lena Palaniyappan

## Abstract

Schizophrenia is characterized by disturbances in both perception and expression that profoundly impair social functioning, yet these domains are typically studied in isolation. Predictive processing theories suggest that such symptoms may arise from abnormal updating of internal models, but the neural mechanisms linking perceptual and expressive dysfunction remain unclear. Using magnetoencephalography during multisensory perception and motor tasks, we tested whether beta-band activity (15-30 Hz), a neural signal implicated in predictive control, provides a shared substrate across domains. Compared to healthy individuals, patients with schizophrenia showed attenuated event-related modulation of beta activity, including weaker beta suppression during sensory processing and delayed or reduced post-movement beta rebound. These abnormalities were associated with a widened audiovisual temporal binding window, indicating atypical multisensory integration. Multivariate analyses further revealed that reduced beta modulation across sensory and motor systems covaried with impoverished semantic diversity, simplified syntactic structure in natural speech, and greater clinical symptom burden. Notably, beta abnormalities emerged as distributed latent patterns spanning sensory, motor, and frontotemporal regions. Together, these findings identify diminished event-related beta modulation as a common neural signature linking disrupted multisensory integration, action monitoring, and language organization in schizophrenia. We situate perceptual and expressive impairments within a unified framework of predictive dysfunction and advance a mechanistically grounded account that highlights beta dynamics as a promising target for future mechanistic and translational studies in psychosis.

**Significance Statement:** Schizophrenia disrupts both how people perceive the world and how they express their thoughts, yet these symptoms are usually studied separately. Using magnetoencephalography during multisensory and motor tasks, we identify a shared neural mechanism linking these domains: reduced event-related modulation of beta-band (15–30 Hz) activity, a signal implicated in predictive control. Compared to healthy individuals, patients showed diminished beta changes during sensory processing and movement completion, suggesting reduced flexibility in updating internal models. Importantly, this neural pattern covaried with abnormal audiovisual binding, disorganized natural speech, and greater clinical severity. These findings reveal distributed beta modulation as a cross-domain neural marker of predictive dysfunction, offering a unified framework for understanding perceptual and expressive disturbances in schizophrenia.

## Introduction

A person with schizophrenia describes that the actors in a Netflix show produce ‘fake speech’, but their real voices come from the radiator vent in his room. When describing this experience, his speech appears disconnected with many ideas weakly developed - he moves from the Netflix ‘TV serial’ to having ‘cereal’ for ‘fastbreak’ to ‘the vent making the room dry’. What links these seemingly disparate symptoms: one perceptual, one expressive? Both perception and language production depend fundamentally on the brain’s ability to generate predictions, compare them against incoming evidence, and flexibly update internal models. Within this predictive processing framework, beta oscillations (15-30 Hz) are a key converging measure of interest that have been proposed to support the maintenance and context-dependent updating of top-down predictive models, including cross-modal integration across the cortex (1–3). Here we show that attenuated modulation of beta-band oscillations, a signal crucial for predictive control of internal models, offers a unified, mechanistically grounded account of both perceptual aberrations and expressive symptoms in psychosis.

Multisensory integration (MSI) allows the brain to combine information from different sensory modalities to form a unified perceptual experience. This process is particularly critical when sensory inputs arrive at different times, requiring the brain to determine whether asynchronous signals originate from the same event. The presence of a temporal binding window (a time interval within which stimuli from different modalities are perceived as synchronous (4)) reflects the brain’s tendency to ascribe common cause to events a priori (i.e., internal predictive model of sensory world (5)). This window is increased in schizophrenia and associated with perceptual disturbances (6) and speech processing (7), and in autism in relation to communication (speech production) deficits (6, 8). A widened temporal binding window indicates an overreliance on the internal model of common cause (5) that is impervious to revisions (9, 10). Alternative accounts, including increased sensory noise, attentional differences, or decision-level biases, could also contribute to broader binding windows in schizophrenia (11–13). Nonetheless, in the present study, the interpretation of reduced predictive flexibility is motivated not by the behavioral effect alone, but by its covariation with event-locked beta modulation as a flexibility index for internal models.

We adopt a specific and constrained interpretation of beta-band dynamics, offering a unified, mechanistically grounded account of both perceptual aberrations and expressive symptoms in psychosis. Rather than testing whether beta oscillations encode the content of predictions, their confidence, or precision-weighting per se, we focus on event-related beta modulation as an index of the flexibility with which internal models are updated in response to task-relevant sensory and motor evidence. Under this formulation, attenuated beta modulation reflects reduced context-dependent manipulation of an ongoing predictive state, consistent with an insufficiently updated internal model. Alternative formulations, such as beta indexing top-down predictions or precision-weighting, are treated here as compatible computational accounts rather than distinct mechanisms directly adjudicated by the present data. In schizophrenia, predictive processing accounts propose aberrant precision weighting between prior beliefs and sensory evidence, resulting in internal models that may be insufficiently updated or overly constrained depending on context and hierarchical level (14–18). Neural processes underlying reduced flexibility in internal prediction models may contribute to both disrupted MSI and speech production abnormalities in schizophrenia.

Three lines of evidence converge to identify the inferior frontal gyrus (IFG) as a critical hub where neural processes underlying predictions may break down in schizophrenia. IFG is critical for multisensory integration, particularly in processing temporal relationships between auditory and visual stimuli. Neuroimaging studies of audiovisual timing judgments and asynchrony detection implicate a right-lateralized fronto-insular and prefrontal network that includes the right IFG, suggesting this region contributes to evaluating and updating perceived audiovisual timing (19–21). Neurophysiological evidence further indicates that neural activity in IFGs reflects supramodal, post-sensory processes related to temporal prediction, integration, and decision-making, typically emerging several hundred milliseconds after stimulus onset rather than during early sensory encoding (22, 23). Meanwhile, converging evidence from lesion, functional Magnetic Resonance Imaging (fMRI), and electrophysiological studies indicates that the right IFG acts as a key control node within a fronto-basal-ganglia network supporting response inhibition and action cancellation, particularly when actions must be withheld or updated based on changing sensory information (23–27). IFG-mediated control is recruited when incoming sensory evidence violates expectations or signals the need to interrupt an ongoing or planned motor response. As such, IFG sits at the interface between sensory evaluation and motor output, enabling flexible adjustment of behavior based on temporally structured multisensory input.

Echoing the right IFG, left IFG involvement is more consistently observed in multisensory tasks that emphasize linguistic, semantic, or categorical processing demands, such as audiovisual speech perception or conflict resolution between competing interpretations (28–32). In particular, the left IFG is also critical for speech production, supporting the controlled selection (33–36), hierarchical sequencing (36–38), and predictive planning of linguistic and motor representations (39, 40). Left IFG also plays a critical role in the two characteristic speech production abnormalities in schizophrenia: reduced lexico-semantic diversity (41, 42) and syntactic complexity (43–46). Together, these findings suggest that impairments in perception, motor control, and speech may reflect disruption of a latent IFG mediated control process rather than independent domain-specific deficits. Critically, IFG contributions may be expressed through the timing, coordination, or control of beta dynamics across distributed sensorimotor networks.

If the IFG serves as an anatomical convergence point, event-related beta modulation may provide the neural mechanism by which predictions are maintained and updated across perception, action, and speech. Beta oscillations have been implicated in both sensory and motor processing, with a distinct role in post-task cortical responses. Post-stimulus beta suppression reflects cortical engagement and processing of incoming sensory information (47–49) via reconfiguration of sensory predictions (40, 50, 51), whereas post-movement beta rebound (PMBR) signals the termination of motor activity and the stabilization of motor predictions (47, 50, 52, 53) following the somatosensory feedback from action execution (50, 52, 54). These post-task responses in the audio/visual/motor cortices represent critical windows for updating or consolidating the internal models of the sensory/motor environment. This context-dependent beta modulation reflects a distributed, system-level mechanism supporting the maintenance, updating, and coordination of internal models across cortex (50, 54), which are crucial for complex acts such as speech production that demands precise temporal coordination of sensory feedback and motor output (55, 56). Identifying latent patterns of coordinated task-related beta dynamics across sensory cortices (V1, A1), motor cortex (M1), and IFG - the higher-order hub for multisensory integration, temporal prediction, and action monitoring - will enable the study of its distributed system-level disruption in schizophrenia.

Here, we test the hypothesis that schizophrenia is characterized by a failure of context-dependent beta modulation, reflecting reduced flexibility in updating internal predictive models, and that this single deficit manifests across multisensory integration, motor control, language organization, and clinical symptom expression.

## Results

### Patients bind sensations across large temporal windows

Auditory and visual stimuli were presented at different onset asynchronies (SOAs see Method). A relative measure of the likelihood to bind perception of obviously asynchronous stimuli was measured as the ratios of average synchronous response rate (SRR) in obvious (e.g., 750, 1000 ms SOAs) versus ambiguous (e.g., 250, 500 ms SOAs) conditions (see Supplementary 1 for model-based psychophysical analysis). SRR were computed for conditions where the preceding trial was auditory-first (AF) and for those where it was visual-first (VF) separately (although combined analysis shows the same pattern of results). These ratios were significantly different between controls and patients for both the audio-first (t(46) = -2.01, p = 0.05) and visual-first conditions (t(46) = -2.43, p = 0.02), with the patients showing a higher preference for reporting synchronous perception in obviously asynchronous conditions compared to HC (Figure 2). Some patients even reported synchronous perception of obviously asynchronous trials in pAF conditions (SRR ratio > 1), confirming prior observations of a wider temporal binding window in schizophrenia (6).

### Patients showed delayed and weaker motor and sensory-related beta modulation

Mean event-related beta power over time windows laid out in *Table 1* were extracted for each participant and compared between groups (patients vs. controls). Patients showed two separate abnormalities on the same effect of impaired predictive flexibility during movement termination: (i) a delay in the onset of post-movement beta rebound in bilateral IFG (0–0.5 s; p < 0.001; Figure 1A), and (ii) a reduction in beta rebound amplitude in bilateral M1 during the established rebound window (0.5-1.0 s; p < 0.01 for both hands; Figure 1B). For sensory processing, visual cortex showed attenuated early beta suppression in right V1 (0-0.2 s post-visual; p = 0.03; Figure 1C). While hypothesis-driven t-tests on predefined auditory time windows did not reach significance, cluster-based permutation testing revealed a significant later cluster (0.2-0.5 s) in right A1 where patients showed weaker beta suppression than controls, indicating that auditory beta suppression deficits in schizophrenia may be more prominent in sustained processing rather than initial encoding (see Supplementary 2).

**Table 1.**
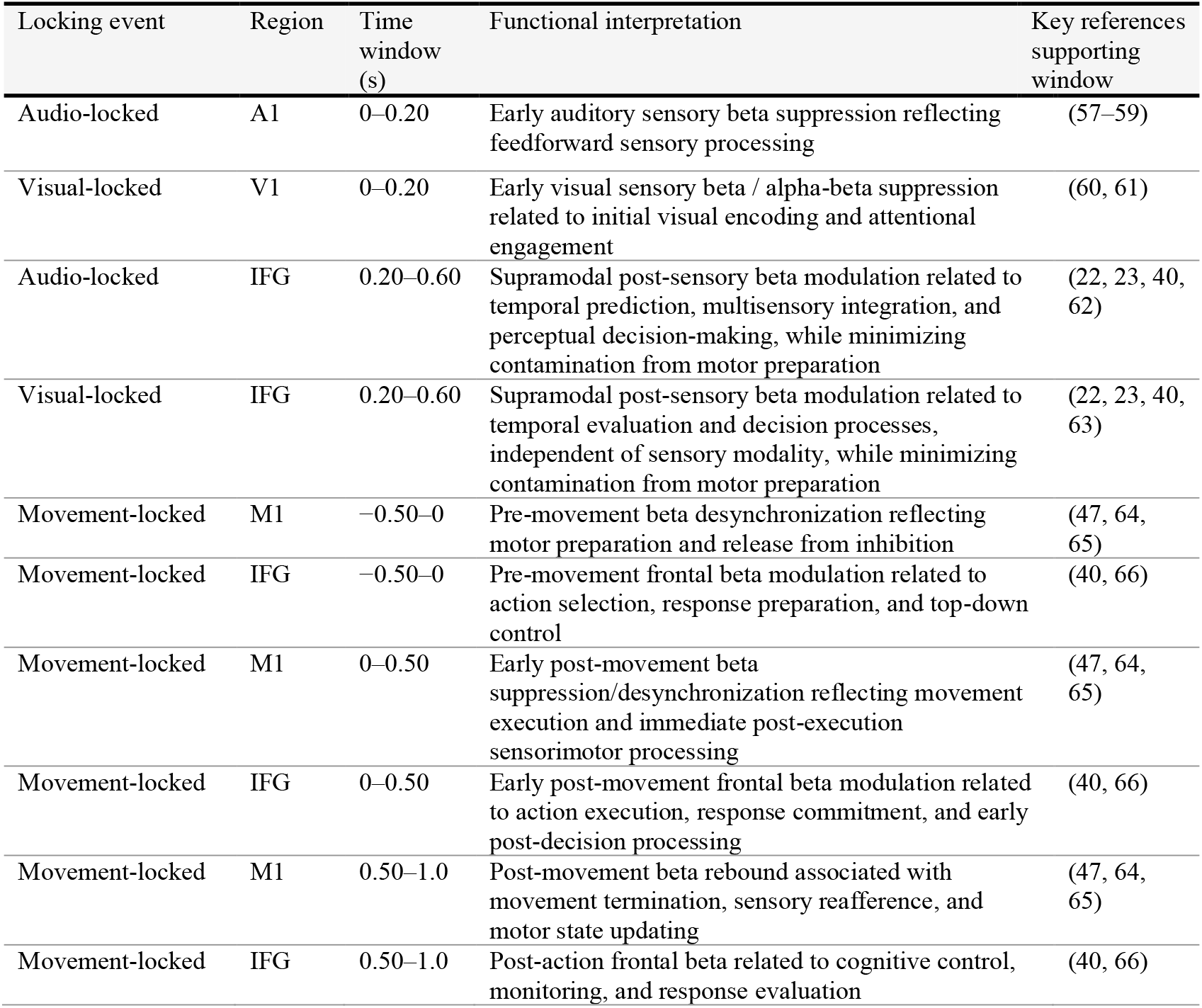
Event-related time windows motivated for beta power extraction.

| Locking event | Region | Time window (s) | Functional interpretation | Key references supporting window |
| --- | --- | --- | --- | --- |
| Audio-locked | A1 | 0–0.20 | Early auditory sensory beta suppression reflecting feedforward sensory processing | (57–59) |
| Visual-locked | V1 | 0–0.20 | Early visual sensory beta / alpha-beta suppression related to initial visual encoding and attentional engagement | (60, 61) |
| Audio-locked | IFG | 0.20–0.60 | Supramodal post-sensory beta modulation related to temporal prediction, multisensory integration, and perceptual decision-making, while minimizing contamination from motor preparation | (22, 23, 40, 62) |
| Visual-locked | IFG | 0.20–0.60 | Supramodal post-sensory beta modulation related to temporal evaluation and decision processes, independent of sensory modality, while minimizing contamination from motor preparation | (22, 23, 40, 63) |
| Movement-locked | M1 | –0.50–0 | Pre-movement beta desynchronization reflecting motor preparation and release from inhibition | (47, 64, 65) |
| Movement-locked | IFG | –0.50–0 | Pre-movement frontal beta modulation related to action selection, response preparation, and top-down control | (40, 66) |
| Movement-locked | M1 | 0–0.50 | Early post-movement beta suppression/desynchronization reflecting movement execution and immediate post-execution sensorimotor processing | (47, 64, 65) |
| Movement-locked | IFG | 0–0.50 | Early post-movement frontal beta modulation related to action execution, response commitment, and early post-decision processing | (40, 66) |
| Movement-locked | M1 | 0.50–1.0 | Post-movement beta rebound associated with movement termination, sensory refference, and motor state updating | (47, 64, 65) |
| Movement-locked | IFG | 0.50–1.0 | Post-action frontal beta related to cognitive control, monitoring, and response evaluation | (40, 66) |

**Figure 1.**
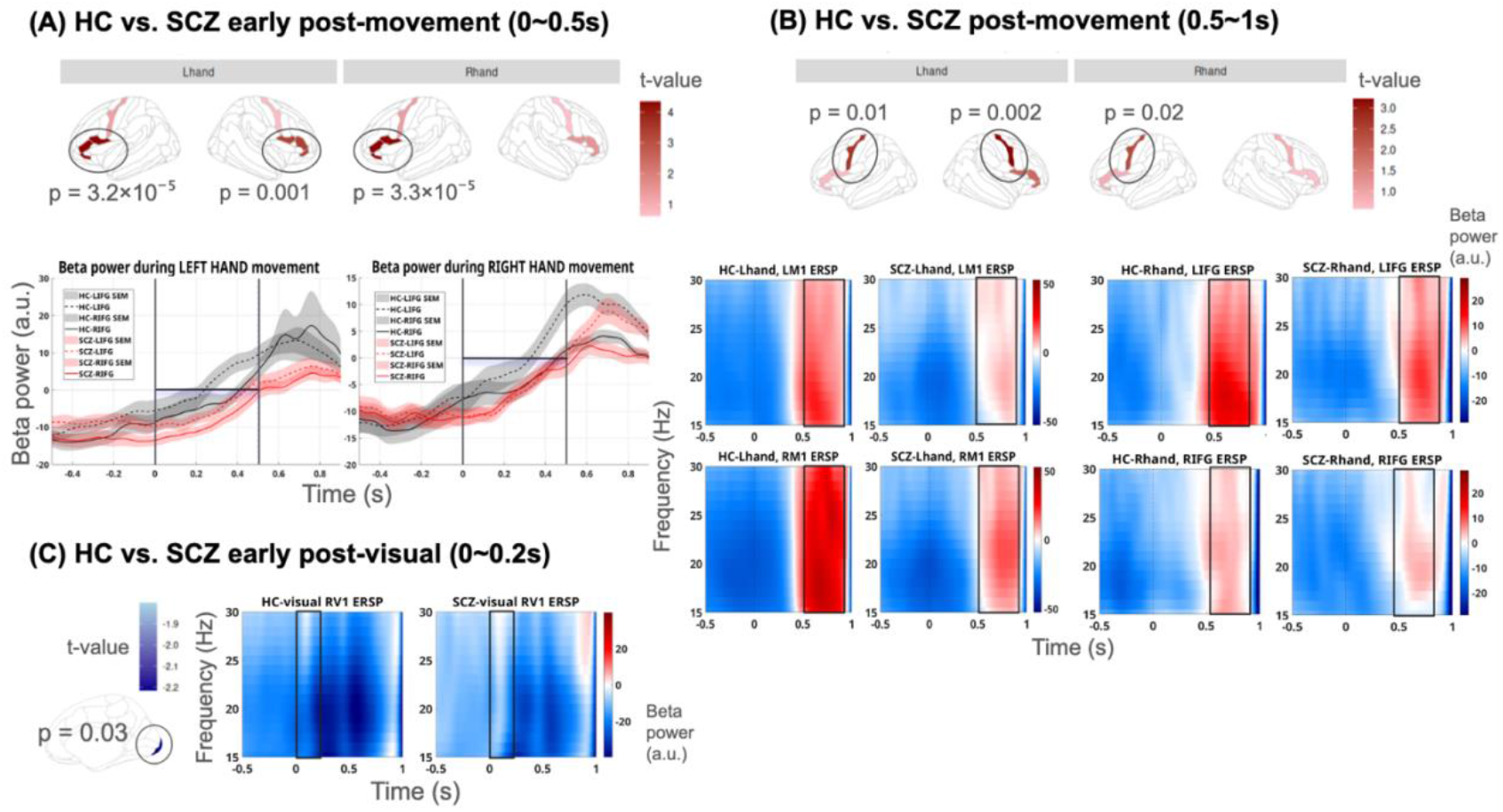
Post-events beta power difference between patients and controls. This figure depicts the post-event beta power difference between patients and controls. Panel A shows t-values comparing mean early post-movement (0∼0.5 s) beta power in M1s and IFGs between controls and patients (top) and the mean beta power with standard error of mean (SEM) sharding across movement-locked timeframe (movement onset = 0 s) (bottom). The black vertical lines in Panel A bottom indicate the timeframe of the significant effect, with the black horizontal line denoting 0 power. Patients had weaker beta power at 0∼0.5 s in IFGs and later beta rebound (beta power increase over 0). Panel B plots t-values comparing mean post-movement (0.5∼1 s) beta power in the M1s and IFGs between controls and patients (top) and the corresponding movement-locked time-frequency maps. Patients had weaker beta rebound power at 0.5∼1 s in the bilateral M1s during both hand movements. Panel C shows a significant t-value comparing mean early post visual (0∼0.2 s) beta power in RV1 between controls and patients (left) and the corresponding time-frequency map. Patients had a weaker early beta decrease in RV1.

These convergent findings support our hypothesis that post-sensory beta suppression and post-movement beta rebound are both reduced in schizophrenia. Mechanistically, these deficits indicate impaired flexibility in updating internal models: attenuated beta suppression constrains the incorporation of new sensory evidence into perceptual predictions, while delayed and weakened beta rebound reflects diminished capacity to stabilize motor predictions and effectively terminate motor programs following somatosensory feedback.

### Multisensory binding relates to weaker event-related beta modulation

To investigate whether beta’s role in predictive model flexibility is supported by the association with multisensory binding behavior, behavioral partial least squares correlation (PLSC) was implemented with permutation testing for latent component significance and bootstrap resampling for feature reliability (see Methods). We identified a single significant latent component linking beta-band activity across distributed cortical regions (cf. Table 1) to behavioral measures of audiovisual simultaneity (LV1: permutation p < 0.01; n=5,000 permutations). This latent component revealed a coherent multivariate association between SRR ratio performance and beta-band dynamics spanning primary motor cortex, inferior frontal gyrus, and early sensory cortices (Figure 2). Bootstrap ratios identified a beta pattern characterized by weaker pre-movement beta suppression in motor and IFG regions, reduced post-sensory beta suppression in auditory and visual cortices and IFG, and attenuated post-movement beta rebound in motor cortex (|BSR| > 1.96, 95% confidence; 5,000 bootstraps). Across participants, stronger SRR ratio, reflecting a greater tendency to bind asynchronous sensory inputs, was associated with reduced modulation of beta activity during both sensory processing and movement termination, as reflected in the association between brain and behavioral latent scores (r(44) = 0.42, p = 0.0036). In other words, a broader temporal binding window was associated with reduced beta modulation during sensory processing and movement termination, consistent with impaired updating of internal sensory-motor predictions.

**Figure 2.**
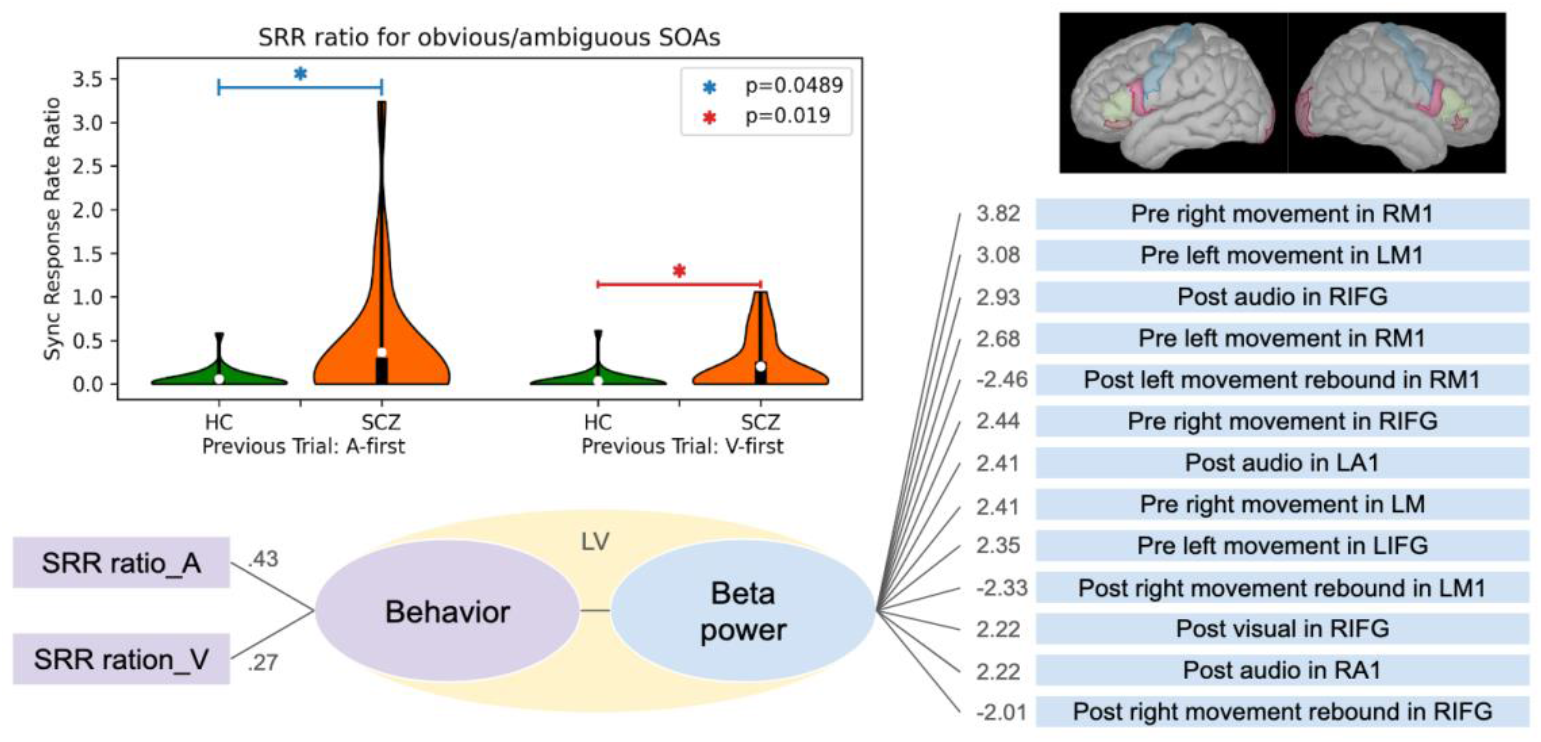
Latent association between beta-band activity and audiovisual simultaneity behavior. Behavioral partial least squares (PLS) identified a single latent component linking beta-band power features with sync response rate (SRR) ratio measures. Top left, violin plots show group differences in SRR ratios for obvious versus ambiguous stimulus onset asynchronies (SOAs), separated by auditory-first (A-first) and visual-first (V-first) preceding trials (p-values indicated). Bottom, numbers adjacent to SRR measures represent *Y loadings* (correlations between each SRR ratio and the behavioral latent score), indicating their contribution to the latent behavioral pattern. Right, beta-band features contributing reliably to the neural latent component are shown with *bootstrap ratios* (salience divided by bootstrap standard error); only features exceeding the bootstrap threshold are displayed. Contributing beta features span motor (M1), inferior frontal gyrus (IFG), and sensory cortices (A1/V1) across pre- and post-event time windows, indicating a distributed beta-band pattern linking sensory-motor processing to audiovisual simultaneity behavior

### Language production abnormalities in patients relate to wide temporal binding window

Natural speech production was recorded and transcribed (see Method) to extract speech features indexing stable patterns of individual speech characteristics instead of a moment-to-moment production dynamics (see Table S1 for language features and Table S2 for NLP models used). To capture patterns of language production abnormalities while managing the high dimensionality of NLP-derived features from natural speech, we performed separate PCAs on semantic and syntactic measures. A semantic component indicating less semantic diversity (i.e., more similar words and sentences with lower semantic density) explained 53.57% of semantic feature variance. A syntactic component indicating lower syntactic complexity (i.e., less complex clauses lacking syntactic depth) accounted for 75.41% of the total syntactic feature variance.

Patients showed semantic impoverishment (i.e., higher scores on the semantic component; t(22.29) = -6.24, p = 0.0000026) and syntactic simplicity (t(42.64) = -4.30, p = 0.000097) compared to controls. Both trained raters blind to the participant groups used the short-form of Thought and Language Index (TLI) scale (67) to assess signs of formal thought disorder, which refers to disturbances in the organization, coherence, and productivity of speech, encompassing both language disorganization (e.g., loosening of associations, peculiar sentences) and impoverishment (e.g., poverty of speech) (68). NLP-derived semantic impoverishment best captured TLI items poverty of speech (t(46) = 4.67, p = 0.000026) and weakening of goal (t(46) = 3.52, p = 0.00099) while the syntactic simplicity best captured peculiar sentences used by the patients (t(46) = 3.29, p = 0.0019).

The tendency to bind sensory percepts over large temporal windows (after removing observations with 0 SRR) was correlated with the degree of semantic impoverishment (SRR ratio_A: r(18) = .53, p = 0.017) and syntactic simplicity (SRR ratio_A: r(18) = .47, p = 0.036; SRR ratio_V: r(12) = .53, p = 0.049), indicating the likelihood of shared mechanisms for both perceptual and expressive behaviours in schizophrenia.

### Attenuated event-related beta modulation covaries with language disturbances

Using behavioral PLSC with permutation testing and bootstrap resampling (see Methods), we examined correlations between a latent beta dynamic reflecting the flexibility of the internal model and the semantic and syntactic language components. For both analyses, permutation testing identified a single significant latent component capturing language-beta covariance (semantic LV1: permutation p < 0.05; syntactic LV1: permutation p < 0.05; n=5,000 permutations).

In the semantic-beta model (Figure 3, top), bootstrap ratios (|BSR| > 1.96) revealed a beta pattern including stronger post-movement beta rebound in bilateral M1s, and stronger post-sensory beta suppression in auditory (LA1) and visual (RV1) cortices, and stronger beta suppression during motor preparation in bilateral M1s. Semantic impoverishment (i.e., higher word-level and sentential semantic similarity with less diversity) was associated with attenuated beta modulation across sensory processing, motor preparation, and movement termination (beta and semantic latent scores, r(46) = .50, p = 0.00029). In the syntactic-beta model (Figure 3, bottom), reliable beta features included reduced post-movement beta rebound in bilateral M1s, decreased post-visual beta suppression in V1s, and weaker beta suppression during motor preparation, particularly in the right M1. Syntactic simplicity (lack of deep, complex ideas) was likewise associated with reduced beta modulation spanning sensory and motor domains (r(46) = .47, p = 0.00079).

**Figure 3.**
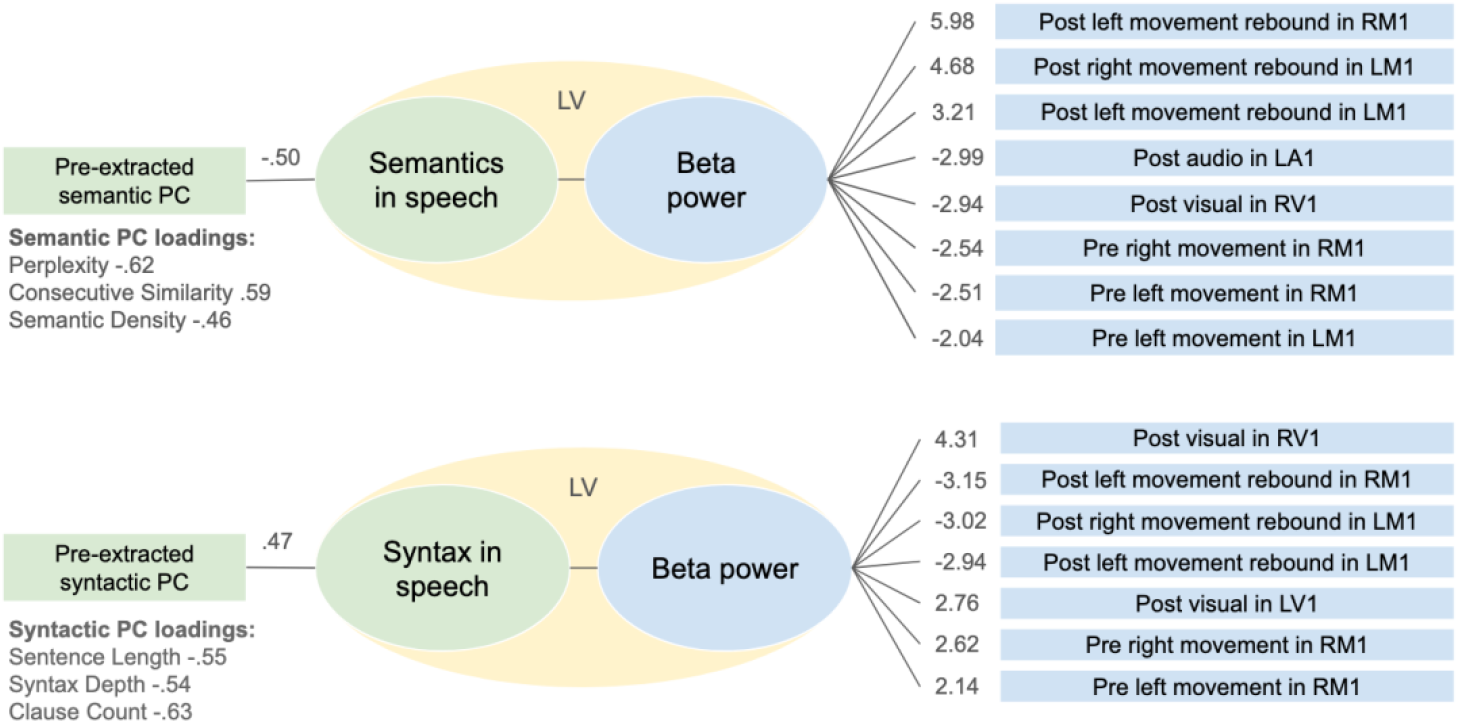
Latent associations between beta-band dynamics and pre-extracted semantic and syntactic language components. Behavioral PLS was used to relate beta-band power features to pre-extracted semantic (top) and syntactic (bottom) principal components (PCs) derived from speech measures. In each analysis, a single latent variable (LV) captured the shared covariance between beta activity and the corresponding language component. Left, schematic representations illustrate the coupling between the language PC and beta power through the LV; PC loadings indicate the contribution of individual speech features to the semantic or syntactic component (only loadings > .40 are shown). Right, beta-band features contributing reliably to each neural latent pattern are shown, with values indicating bootstrap-normalized weights (bootstrap ratios); only features exceeding the bootstrap reliability threshold (|BSR| > 1.96) are displayed. For both semantic and syntactic models, reliable beta features were observed predominantly during pre-movement and post-movement intervals in primary motor cortex, as well as post-sensory responses in auditory and visual cortices, indicating that beta dynamics during sensorimotor processing covary with higher-order linguistic organization.

Together, these results indicate that the disrupted beta dynamics across sensory and motor domains (i.e., weaker beta decrease during sensory processing and motor preparation and weaker post movement beta rebound), which also relate to temporal binding during multisensory integration, covary with stationary language organization features in schizophrenia. Although we did not test for a direct causal pathway, these findings support a shared latent beta abnormality linking disrupted perception, action, and language in schizophrenia.

### Weaker beta modulation of internal model relates to worse psychotic symptoms in patients

With the smaller patient sample size (n = 23) that limits the permutation and bootstrapping methods, we undertook an exploratory assessment of whether the linguistic organization-related beta dynamics also related to clinical severity. One principal component was extracted from 10 clinical scores, explaining 58% of total variance in illness burden (see Table S3). The illness burden was weighted by the core psychotic symptoms (e.g., higher positive and negative symptom severity and lower social functioning measures). Antipsychotic exposure (Defined Daily Dose) did not correlate with illness burden (r(21) = .18, p = .42).

The beta latent scores derived from the semantic-beta and syntactic-beta PLS models were correlated with illness burden: the latent beta dynamics related to semantic impoverishment (Figure 3, top) significantly correlated with illness burden (r(21) = -.46, p = 0.027; weaker semantic-beta modulation seen with more severe burden) while syntactic simplicity related beta (Figure 3, bottom) had a trend level association (r(21) = .39, p = 0.06).

Together, these findings show that beta modulation in schizophrenia is not limited to a single cortical region or task, but emerges as a distributed latent pattern linking sensory processing, motor control, language production, and clinical symptom expression. This consistent signature across different levels of analysis suggests that beta dynamics is a key neural mechanism underlying dysfunctional internal models in psychosis.

## Discussion

We identify attenuated beta modulation as a shared neural signature unifying perceptual, motor, and linguistic dysfunction in schizophrenia. Patients showed reduced beta suppression during sensory processing and delayed, weakened beta rebound following movement. These dynamics predicted multisensory binding deficits, impoverished speech, and illness burden, establishing a mechanistic link across the domains of perception and expression in psychosis.

A central contribution of this work is the specification of how predictive-processing abnormalities manifest in beta dynamics across distinct task contexts. Within predictive coding frameworks, psychosis has been characterized as reflecting either overly strong priors that dominate perception or weak priors that fail to stabilize inference (15–17). Our results refine these accounts by demonstrating that the critical abnormality lies in the dynamics of updating predictive models (indexed by event-related beta modulation) rather than in the absolute strength of priors themselves, consistent with proposals that beta oscillations index the maintenance and precision-weighting of internal models (1, 2, 40). Under this formulation, elevated baseline or resting beta activity, as often observed in schizophrenia (69, 70), can coexist with diminished event-related beta suppression or rebound, because the core deficit lies in its context-sensitive regulation, not oscillatory capacity itself (see (71) for a converging framework).

Consistent with this prediction, we observed insufficient beta suppression during sensory processing, alongside behavioral evidence of a wide binding window. As described above, transient beta suppression reflects a release of top-down constraints that permits sensory evidence to update perceptual predictions (22, 40). Here, the reduced depth of beta suppression in patients indicates that this release is incomplete, consistent with an internal model that resists updating (14, 17). Importantly, this effect was temporally specific and event-locked, rather than a generalized reduction in beta power, supporting the interpretation of impaired modulation rather than diminished oscillatory capacity. The same failure of event-related modulation was expressed during motor behavior as delayed and attenuated post-movement beta rebound. Consistent with the established role of post-movement beta rebound in stabilizing predictive motor states (47, 50, 54), the delayed and weakened rebound observed in patients here suggests a failure to efficiently transition between predictive regimes following action (contrary to the model of imprecise priors (72)).

Within the predictive framework outlined above, the present findings constrain how failures of beta modulation manifest across sensory and motor domains. If beta supports the maintenance of internal models, then reduced flexibility in model updating (14, 15) should manifest as diminished beta modulation following both sensory events that require updating perceptual predictions and motor feedback that signals the need to transition into a post-movement state. Together, these findings demonstrate that a single deficit in predictive flexibility can yield distinct but principled beta abnormalities across task contexts, without invoking multiple mechanisms.

Importantly, these beta abnormalities extended beyond perception and action to natural language production, a domain that critically depends on predictive control. Although we did not measure beta activity during speech production itself, we observed that the same patterns of attenuated sensory and motor beta modulation covaried with reduced semantic diversity and syntactic complexity in spontaneous speech. This cross-domain association supports, but does not by itself establish, the interpretation that beta-mediated predictive mechanisms engaged during sensorimotor processing are also relevant for language generation. This interpretation aligns with converging evidence that beta oscillations support the maintenance and sequencing of linguistic context and the pre-activation of lexical-syntactic representations during language processing, though not directly during language production (39, 73, 74). Under this account, insufficient beta modulation may reflect a failure to flexibly pre-activate and update linguistic representations, biasing production toward high-probability, generic lexical items and simplified syntactic structures, as observed in formal thought disorder (75, 76).

IFG showed temporal delays in beta rebound, echoing its role in coordinating transitions between predictive states rather than expressing large local beta-power changes. It was also involved in multisensory binding associations, consistent with its role in supramodal timing. Although IFG beta did not show robust amplitude reductions or contribute to speech-related covariance, this may reflect that IFG beta is most engaged during active linguistic planning not probed here, or that frontal contributions operate through connectivity rather than local power. The absence of strong IFG beta-speech associations should be interpreted as a constraint on when frontal predictive mechanisms are expressed in oscillatory dynamics, not as evidence against IFG involvement in language.

Speech production relies critically on the integration of auditory feedback and motor planning, and beta dynamics in auditory and motor cortices are well positioned to support the updating of these sensorimotor predictions. Under this interpretation, abnormalities in sensorimotor beta modulation may predict the fidelity of internal models that are subsequently recruited during language generation, yielding impoverished semantic diversity and simplified syntactic structure without requiring direct frontal beta dysfunction. Nonetheless, several factors may explain the limited IFG contribution to speech-related beta covariance in the present data. First, beta oscillations in IFG may be most strongly engaged during active linguistic planning and selection, processes not directly probed by the sensorimotor tasks used here. Second, IFG involvement in language may be expressed through transient, context-dependent dynamics that are less stable across individuals and therefore less likely to emerge in group-level or multivariate covariance analyses. Third, IFG contributions to language may be mediated through connectivity or cross-frequency interactions rather than local beta power, which were not assessed in the present study. Thus, the absence of a strong IFG beta-speech association should not be taken as evidence against IFG involvement in language, but rather as a constraint on when and how frontal predictive-control mechanisms are expressed in oscillatory dynamics.

Limitations include modest sample size, cross-sectional design precluding causal inference, analysis restricted to beta power rather than phase or connectivity, and speech recorded offline rather than during scanning. However, the convergence of effects across behavioral, neural, linguistic, and clinical domains partially mitigates concerns related to sample size, particularly given the use of latent-variable approaches designed to capture shared variance across distributed features rather than isolated effects. Beta bursts and network coordination may offer more direct insight into predictive signaling (77–79). Medication effects, while not correlated with clinical scores, cannot be fully ruled out. Future studies should examine first-episode patients, employ longitudinal designs, and directly measure oscillatory activity during language production.

In summary, by demonstrating that attenuated event-related beta modulation relates to aberrant multisensory integration, disorganized speech, and clinical severity, this work advances a unified account of perceptual and expressive dysfunction in psychosis. More broadly, it identifies distributed beta modulation as a cross-domain neural phenotype of predictive dysfunction, a system-level signature of reduced flexibility in updating internal models across perception and action. By linking this phenotype to behavioral and linguistic disturbances, our findings position beta dynamics not only as a biomarker of illness burden in schizophrenia, but as a mechanistically grounded target for interventions (see (80) for a recent review on event-related beta as neuromodulatory treatment target) aimed at restoring predictive flexibility in disorders of disrupted inference.

## Materials and Methods

### Participants

This study followed a case-control design involving 25 patients with schizophrenia (age: 33.26 ± 9.03; 8 females) and 25 age and sex matched healthy controls (age: 33.36 ± 9.26; sex: 7 females) (education unmatched but controlled for in analysis; t(45.98) = 6.23, p = 0.00000013). The study was approved by the Research Ethics Boards of McGill University and the Douglas Mental Health University Institute, and all participants provided written informed consent in accordance with institutional guidelines. Inclusion criteria included English or French-speaking participants, male or female, aged 18-65 years. Patients were recruited from the inpatient, outpatient, and community services caseload held by consultant psychiatrists working at McGill-affiliated hospitals in Montreal, Quebec. Patients had to be previously diagnosed with schizophrenia or schizoaffective disorder based on the Diagnostic and Statistical Manual of Mental Disorders 5th Edition (DSM 5). Consenting participants eligible for the study underwent symptom and speech assessment, as well as MEG and MRI recordings. Two patients were excluded from MEG analysis (MEG N=48) due to dental and piercing-related artifacts. A different set of two patients (behavioral N=48) were excluded from behavioral analysis due to inability to follow task instruction. When associating MEG with behavioral data, the sample size was 46.

Clinical assessments including SOFAS (81), PANSS-8 (82), and CGI-S (83) were implemented. A total of 10 numeric clinical rating scores were measured. To extract a single latent factor indicative of illness burden before associating clinical scores with MEG and speech metrics, principal component analysis (PCA) was used to extract the first component (explaining 58% of total variance) (see Supplementary Table S3). For each participant, a defined daily dose (DDD) was determined for psychotropic medications (antipsychotics, antidepressants, benzodiazepines, stimulants, anticonvulsants, beta blockers, and anticholinergic drugs separately). Participants provided a list of current medications, including maximum dose, and duration. Where necessary, medication dosage was verified in their medical file. DDD per drug class was determined by ATC/DDD Index (https://atcddd.fhi.no/atc_ddd_index/) in which the dose the maximum dose the participant was taking was divided by the DDD.

### Multisensory integration task

During MEG recording, participants completed an audio–visual simultaneity-judgment task. Each trial presented a white square (visual; V for 50-ms) at screen center and a 1000-Hz tone (auditory; A for 50-ms) with one of 11 stimulus onset asynchronies (SOAs): 0 ms, ±100 ms, ±150 ms, ±200 ms, ±750 ms, and ±1000 ms (negative values = audio first). Most SOAs were presented 8 times per block for each lag direction (e.g., +100 ms and –100 ms each repeated 8 times). The ±750-ms and ±1000-ms SOAs were presented twice per block and served only as “obviously asynchronous” anchors.

Visual stimuli were back-projected at 120 Hz, and tones were delivered binaurally via foam-tip earphones. On each trial, participants judged the pair as synchronous (S) or asynchronous (A) and responded using button pads positioned on both sides of the MEG chair. The S/A mapping to left/right buttons was pseudo-randomized within each block to counterbalance response laterality. After stimulus presentation, the letters “A” and “S” briefly appeared on the corresponding response sides to minimize rule-forgetting; they disappeared immediately upon button press. The task comprised five blocks of 64 randomized trials each (total: 350 trials). Blocks began when participants pressed the button assigned to the synchronous response. Inter-trial intervals were jittered between 2.3 and 2.8 s, during which only a fixation cross was displayed.Two brief preparation blocks preceded the main task. The first familiarization block presented each SOA once in a fixed sweep order (–1000 ms to +1000 ms), with the SOA value shown beforehand to give participants an intuitive reference. The second was a short practice block presenting each SOA once in random order using the standard trial format.

### Behavioral analysis

For each participant, trials were pooled into conditions according to three properties: 1) the magnitude of stimulus onset asynchrony (SOA) value in millisecond, 2) the sign of the SOA, which corresponds to the leading sensory modality of the stimulus (negative being auditory-first and positive being visual-first), and 3) the leading sensory modality of the stimulus from the previous trial. For each condition we computed a synchronous response rate (SRR), which corresponds to the percentage of trials in a given condition for which the participant reported perceiving the stimulus as synchronous. We categorized conditions with stimulus onset asynchrony (SOA) of ±750ms and ±1000ms as “obvious” and those with SOA of ±100ms, ±150ms, and ±200ms as “ambiguous”. We averaged the SRR for obvious and ambiguous conditions before computing the ratio between these average SRR in conditions where the trials were following an auditory-first trial (pVA) and in those where they were following a visual-first trial (pVF), resulting in 2 ratios per participant. We compared the pVA and pVF ratios between groups using two-sampled independent t-tests with a p=0.05 significance threshold. Note: Two participants in the patient group were removed due to clear issues in their response patterns (e.g., giving the same response for entire blocks; N=48). See Supplementary 1 for model-based psychophysical analysis on subjective simultaneity and temporal recalibration.

### Natural speech recording and analysis

Speech samples were collected using the Discourse in Psychosis Consortium protocol (Discourse in Psychosis Consortium, 2021). The speech protocol took around 20 minutes of administration and consisted of 7 sections, including free speech, free personal narrative, health narrative, picture description, storytelling, dream reports, reading and recall. Audio recordings from participants’ interviews are transcribed to text using WhisperX (https://github.com/m-bain/whisperX) before manual correction of transcription and speaker diarization was done. All sentences from the interviewer were removed such that the textual data submitted to the analysis pipeline only contains utterances from the participant. The text associated with each participant was analyzed independently to generate a set of 30 speech features describing the language characteristics of the participant. The speech features extraction pipeline relies on both basic NLP algorithms and various types of LLMs (see Table S2). From our speech samples, semantic and syntactic features were extracted. Semantic features capture how meaning unfolds across discourse, using measures such as consecutive sentence similarity (average cosine similarity between sentence embeddings), lexical predictability (perplexity and pseudo-perplexity), and semantic density (the proportion of meaningful components within a sentence vector). Syntactic features index structural complexity based on part-of-speech parsing, including sentence length, syntactic tree depth, and clause count.

### MEG Recording

MEG data were acquired using a whole-head 275-channel CTF MEG system (CTF/VSM, Coquitlam, Canada) housed in a magnetically shielded room at the Montreal Neurological Institute. Signals were recorded with axial gradiometers and online third-order synthetic gradiometer noise reduction, sampled at 1200 Hz. Participants were seated upright with head support to minimize movement. Head position was monitored continuously using three localization coils placed at the nasion and left/right preauricular points and re-localized at the start of each block. Individual head shapes (∼100 scalp points including fiducials) were digitized using a FASTRACK electromagnetic tracking system (Polhemus, USA) and used to co-register MEG data to each participant’s structural MRI via rigid-body transformation optimized by dense scalp surface alignment. Horizontal and vertical electrooculography (EOG) and electrocardiography (ECG) were recorded concurrently to aid artifact identification. Datasets with within-block head movement exceeding 5 mm or transient displacements >10 mm were flagged for review. Noisy or malfunctioning channels were identified using CTF system diagnostics and excluded prior to preprocessing.

### Structural MRI Acquisition for MEG Co-registration

T1-weighted images were collected from all participants with the 3T Siemens Prisma MRI scanner at the Neuro. The cortical surface used in MEG source modeling was derived using freesurfer 7.4.1 (recon-all) on a preprocessed skull-stripped image obtained by applying presurfer (ref to presurfer: https://github.com/srikash/presurfer) on the UNI and INV2 maps obtained from the MP2RAGE sequencing. The resulting cortical surface was then imported into Brainstorm to perform, and refine, co-registration between the MRI and MEG data by aligning the Polhemus FASTRACK fiducials (75-100 headpoints including nasion and preauricular points) to their corresponding MRI landmarks, followed by refinement using the densely digitized scalp surface points. The resulting transformation matrix was used for forward model computation and subsequent source-space analyses.

### MEG preprocessing and time-frequency analysis

MEG preprocessing and source modeling were performed in Brainstorm using automated MATLAB scripts. Continuous CTF recordings and corresponding empty-room data were imported, realigned to digitized head points, and band-pass filtered (0.1–100 Hz). Auditory onsets were identified from the analog trigger channel. Cardiac and ocular artifacts were detected using peak-based methods on ECG and VEOG channels; signal-space projection (SSP) components were computed, manually reviewed, and applied. Cleaned data were epoched from –3 to 1 s relative to left/right button presses, auditory onsets, and visual onsets.

Noise covariance was estimated from empty-room recordings and data covariance from all trials. Forward models were constructed using overlapping spheres on individual MRIs, and source estimation employed an LCMV beamformer with neural activity index normalization and unconstrained dipoles. Dipole orientations were reduced using PCA (first component) computed over a baseline window (–2.5 to –2.0 s). Source estimates were averaged across runs per subject and projected onto a standard cortical template.

Regions of interest (ROIs) were defined using the Destrieux atlas (84), including inferior frontal gyrus subdivisions, primary motor cortex (precentral gyrus/M1), primary visual cortex (occipital pole/V1), and primary auditory cortex (transverse temporal sulcus/A1), with all vertices included. Time–frequency decomposition was performed on source-level ROI signals using Morlet wavelets (1–60 Hz, 1-Hz steps), yielding baseline-normalized event-related spectral perturbations (ERSP). Motor-related activity was baselined to –2.5 to –2.0 s to avoid movement preparation, while sensory-related activity was baselined to –1.5 to –1.0 s separately for auditory-first and visual-first trials to avoid stimulus contamination. These ERSP maps constituted the measures used for group-level analyses.

### Beta-band power extraction and time-window selection

To obtain subject-specific indices of beta-band modulation, for each subject, run-averaged ERSPs were computed for motor (left/right button press) and sensory (auditory, visual) events. Within anatomically defined regions of interest (ROIs), we computed the average ERSP map over all vertices. Average beta (15-30 Hz) powers were then extracted from each ROI’s ERSP map according to the event and time-window specified in Table 1.

We focused on beta-band power modulation as a robust, interpretable first-order index of predictive control dynamics, noting that burst-based, phase-based, or connectivity analyses represent complementary levels of description beyond the scope of the present study.

### Statistical analysis

For beta power-group difference from hypothesis-driven time windows specified a priori (Table 1), we report uncorrected Welch’s two-sample t-tests at alpha=0.05. For exploratory time-frequency analyses not driven by specific hypotheses, we employed cluster-based permutation testing (5,000 permutations) to control family-wise error rate (see Supplementary 2).

To relate beta-band features to behavioral, speech, and clinical measures, we used behavioral partial least squares (PLS) as implemented in the pyls Python package. For each analysis, we constructed an X matrix of subject-wise beta power features and a Y matrix of either behavioral (audiovisual simultaneity) or speech (semantic/syntactic). All variables in X and Y were z-scored across participants prior to analysis. Behavioral PLS decomposes the cross-block covariance between X and Y into orthogonal latent variables (LVs) that maximally capture shared covariance. Statistical significance of each LV was assessed using permutation testing (5,000 permutations of the subject labels), and the stability of individual feature contributions was evaluated using bootstrap resampling (5,000 bootstraps) to obtain bootstrap ratios (salience divided by bootstrap standard error). Features with |bootstrap ratio| > 1.96 were interpreted as reliable contributors to a given LV.

Pearson correlations were used to reveal association between clinical component and beta latent scores.

## Acknowledgments and Funding

This research is supported by Mitacs Accelerate Fellowship (IT44173) to HTW, McConnell Brain Imaging Centre (Magnetoencephalography in Psychosis to SB and LP), and Quebec Bioimaging Network (QBIN 35450 grant) to LP

## Data sharing

Code and material will be made available on Open Science Framework upon publication. Temporary link: https://osf.io/hjfz4/overview?view_only=6d2d6dbd1a9848648a2b010fed96aea2. The datasets generated during and/or analyzed during the current study are available from the corresponding author on request. Please contact Dr. Lena Palaniyappan at.

