## Supplementary for "Distributed neurophysiological dynamics link perception, action, and language in schizophrenia"

**This PDF file includes:**

Supporting text
Figures S1 to S2
Tables S1 to S3
SI References

### Supporting Information Text

#### 1. Psychophysical behavioral analysis/results

**Methods.** The pooling of trials into conditions yielded a total of 22 conditions with an associated response rate values per participant: 11 SOA and 2 categories of trial sequencing (pAF and pVF). The SRR values for each category are used to derive psychometric curves by fitting the mean, standard deviation, and amplitude of a Gaussian model to the response rate values (in %). The Gaussian model's mean parameter corresponds to the point of subjective simultaneity for pAF (Point of Subjective Simultaneity (PSS) A) and pVF (PSS V) conditions. Temporal recalibration (TR), estimated as the difference between PSS A and PSS V values, reflects rapid adjustments in perceptual processing made to align asynchronous audio-visual sensory inputs together and facilitate multi-modal sensory integration (1).

We derived the parameters of Gaussian models for each subject in both pAF and pVF conditions alongside their associated Gaussian curves using all SOA values and repeated the process with only ambiguous conditions. In all cases, participants who estimated TR magnitude exceeded the realistic range (±300ms) or for whom the curve fitting algorithm did not converge for either pAF or pVF conditions were excluded from analyses involving the associated PSS and TR values. For visualization purposes, we also derived group-average PSS and TR values by averaging the SSR of all participants that were not rejected in each group and fitting a Gaussian model to the resulting SSR values. We tested the significance of TR values in each group (one-sample *t*-test against 0).

**Results.** When using all SOA conditions, no control was excluded, and 6 patients were excluded due to non-converging curve fitting or TR values exceeding the realistic range. The TR values for HC were significant (one-sample *t* test against 0, $t(24)=4.237$, $p=0.0003$) with an average of 30ms, replicating results from previous studies (Supplemental Figure 1, top right). The TR values for the patient group were also significant (one-sample *t* test against 0, $t(22)=2.484$, $p=0.0231$), with an average of 41ms (Figure S1, bottom right). No significant differences were found between groups for any of the PSS or TR measures using two-sample independent *t* tests (Welch's two-sample t-test, PSS_VF: $t(44.89) = 1.55$, $p = .13$; PSS_AF: $t(42.90) = .95$, $p = .35$; TR: $t(30.62) = 1.16$, $p = .26$).

When using only ambiguous SOA conditions, 1 HC participant was excluded and 4 patients were excluded due to non-converging curve fitting or TR values exceeding realistic range. The TR values for HC remained significant (one-sample *t* test against 0, $t=4.124$, $p=0.0004$) with a slightly lower average of 22ms (Supplemental Figure 2, top right). The TR values for the patient group were no longer significant (one-sample *t* test against 0, $t=1.20$, $p=0.2365$), with an average of 22ms (Figure S2, bottom right). No significant differences were found between groups for any of the PSS or TR measures using two-sample independent *t* tests.

#### 2. Cluster-based permutation test on ERSP maps between patients and controls

**Method.** To assess group differences in event-related spectral perturbations (ERSPs) without imposing a priori constraints on time-frequency windows, we performed non-parametric cluster-based permutation tests using FieldTrip. Source-localized ERSP maps were extracted from individualized peak vertices within bilateral primary motor cortex (M1), inferior frontal gyrus (IFG), primary auditory cortex (A1), and primary visual cortex (V1) for motor, auditory, and visual events, respectively. ERSPs were computed across frequencies spanning 15-30 Hz and time windows from -0.5 to 1 s relative to event onset. Prior to statistical testing, the time dimension was down-sampled to ~20-ms bins to improve computational efficiency while preserving temporal resolution.

Group differences between healthy controls (HC; $n = 25$) and patients with schizophrenia (SCZ; $n = 23$) were tested using independent-samples t-statistics with cluster-based correction for multiple comparisons across time and frequency. Clusters were formed by identifying adjacent time–frequency samples exceeding a cluster-forming threshold of $p < 0.05$ (two-sided). Cluster-level statistics were computed using the maximum cluster mass (sum of t-values), and significance was assessed via Monte Carlo permutation with 1,000 randomizations. The family-wise error rate was controlled at $\alpha = 0.05$ (two-tailed). This approach provides robust control over multiple comparisons while remaining sensitive to spatially and temporally extended effects.

**Result.** Cluster-based permutation testing on time–frequency representations (15-30 Hz, -0.5 to 1 s) identified significant group differences in beta-band activity for both motor- and sensory-locked events (*Figure S2*). For motor events, significant clusters were observed in primary motor cortex contralateral to the moving hand (RM1 for left-hand movements, LM1 for right-hand movements) and in inferior frontal gyrus (LIFG and RIFG). These clusters were temporally extended over the peri- and post-movement interval (≈0~1 s) and indicated delayed and attenuated post-movement beta rebound in patients relative to controls. For sensory events, significant beta clusters were found in right primary auditory cortex (RA1) following auditory onsets (≈0~0.5 s) and in right primary visual cortex (RV1) from pre-visual to around visual onsets (-0.5~0.2s). These sensory clusters reflected weaker event-related beta suppression in patients. Although cluster-based tests are conservative and favor temporally and spectrally extended effects (2, 3), the presence of significant clusters in both motor and sensory cortices indicates that the observed beta abnormalities are robust at the group level, with motor-related differences showing the broadest temporal extent.

**Supplementary Information Figures**

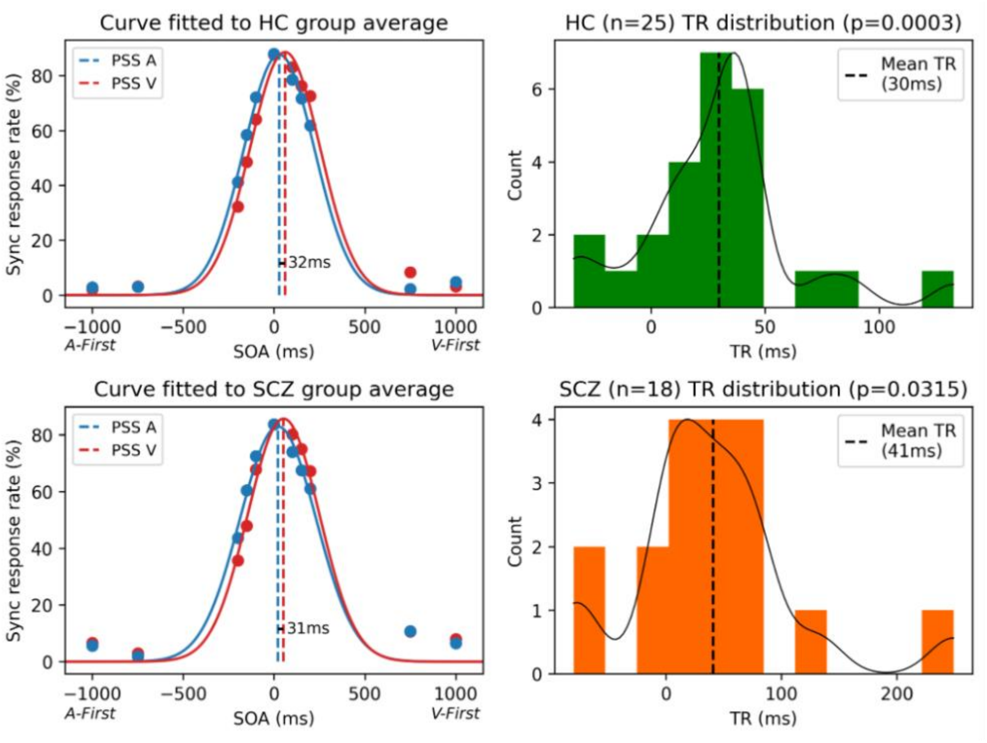


**Figure S1. Summary of temporal recalibration analysis using all SOA conditions for both groups**.
The left column illustrates the gaussian models (solid lines) fitted to HC (top) and SCZ (bottom) group-averaged SSR values, with their respective PSS (dashed lines) alongside the associated temporal recalibration (black line). The right column illustrates the distribution of temporal recalibration values (histogram) for HC (top) and SCZ (bottom) alongside a matching kernel density estimate (solid black line) and the average TR (dashed line).

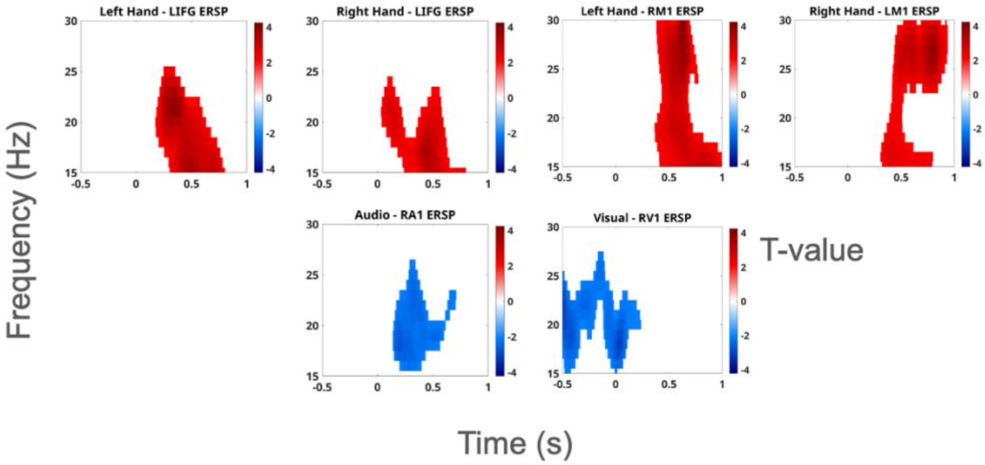


**Figure S2. Cluster-based permutation results for beta-band activity.**
Significant time–frequency clusters (15–30 Hz) identified by cluster-based permutation testing comparing patients with schizophrenia and healthy controls are shown for motor-locked (left and right hand movements) and sensory-locked (auditory and visual onsets) events. Color values represent t-statistics for group differences, with red indicating higher beta power in controls than patients and blue indicating lower beta power in controls than patients. For motor events, significant clusters were observed in primary motor cortex (RM1 and LM1) and inferior frontal gyrus (LIFG and RIFG), spanning peri- and post-movement intervals and reflecting delayed and attenuated post-movement beta rebound in patients. For sensory events, significant clusters were observed in right primary auditory cortex (RA1) and right primary visual cortex (RV1), primarily during early post-stimulus intervals, reflecting weaker event-related beta suppression in patients. Axes denote time relative to event onset (s) and frequency (Hz).

**Supplementary Information Tables**

**Table S1. Language PC loadings and speech feature definitions**

| **Speech Domain** | **Feature Name** | **Loading on PC** | **Feature Description** |
|---|---|---|---|
| **Semantics** | Consecutive Similarity | **.59** | Sentence-level cosine similarity. Higher similarity score indicates greater similarity between neighboring sentences in the embedding space. |
| | Perplexity | **-.62** | Perplexity of a generative model when evaluating the text from a participant. Higher perplexity reflects a lower capacity of the model to predict the subject's word choice. |
| | Pseudo Perplexity | .23 | Pseudo-perplexity represents the ability of a mask filling model to guess a content token. Higher score is more unusual word choice given the preceding and following context |
| | Semantic Density | **-.46** | Ratio of meaningful to content words in a sentence, where meaningful components are derived via vector unpacking, applying gradient descent to word2vec sentence embeddings to approximate the original semantic vector. A higher value shows higher semantic density. |
| **Syntax** | Sentence Length | **-.55** | Number of words in a sentence. |
| | Syntax Depth | **-.54** | Depth of the syntax tree obtained by parsing a sentence through part-of-speech tagging. A higher value indicates a complex syntax. |
| | Clause Count | **-.63** | Number of clauses as recognized by pattern matching on the part-of-speech tags. |

Note. Feature loadings above .4 are bolded.

**Table S2. NLP/LLM models for speech feature extraction**

| Model Type | Description | Metrics Relying on the Model |
|---|---|---|
| **Causal Language Model (Decoder-Only)** | Decoder-only transformer model with causal attention layers to produce unidirectional attention. It is used to generate vectors representing logits for the next token prediction. | Perplexity |
| **Mask-Filling Model** | Bidirectional embedding model with a mask prediction head to produce an output vector representing the logits corresponding to the masked token. | Pseudo Perplexity |
| **Word2Vec** | Context-independent word-level embedding model composed of a large dictionary of vectors. It maps each word to a vector corresponding to its meaning. The sentence level embedding is calculated by averaging across word vectors. | Semantic Density |
| **Does not Rely on a Model** | NA | Sentence Length |
| | | Syntax Depth |
| | | Clause Count |

**Table S3. Clinical measures and their loadings on the clinical PC**

| **Clinical scores** | **Clinical measure** | **1st component loadings** |
|---|---|---|
| SOFAS | Social and Occupational Functioning Assessment Scale | **-.34** |
| PANSS-P1 | Delusion | **.36** |
| PANSS-P2 | Conceptual disorganization | .26 |
| PANSS-P3 | Hallucinatory Behavior | .30 |
| PANSS-N1 | Blunted affect | **.33** |
| PANSS-N4 | Passive/apathetic social withdrawal | **.37** |
| PANSS-N6 | Lack of spontaneity and flow of conversation | .17 |
| PANSS-G5 | Mannerisms/Posturing | .17 |
| PANSS-G9 | Unusual thought content | **.36** |
| CGI-S | Considering your total clinical experience with patients with schizophrenia, how severely ill has the patient been during the course of illness? | **.40** |

Note: Feature loadings above .3 contribution to the component are bolded.

**Supplementary Information References**

1. T. Lennert, S. Samiee, S. Baillet, Coupled oscillations enable rapid temporal recalibration to audiovisual asynchrony. *Commun Biol* **4**, 559 (2021).

2. E. Maris, J.-M. Schoffelen, P. Fries, Nonparametric statistical testing of coherence differences. *Journal of Neuroscience Methods* **163**, 161–175 (2007).

3. J. Sassenhagen, D. Draschkow, Cluster-based permutation tests of MEG/EEG data do not establish significance of effect latency or location. *Psychophysiology* **56**, e13335 (2019).
